# EDEM3 modulates plasma triglyceride level through its regulation of LRP1 expression

**DOI:** 10.1101/742353

**Authors:** Yu-Xin Xu, Gina M. Peloso, Taylor H. Nagai, Taiji Mizoguchi, Amy Deik, Kevin Bullock, Honghuang Lin, Kiran Musunuru, Qiong Yang, Ramachandran S. Vasan, Robert E. Gerszten, Clary B. Clish, Daniel Rader, Sekar Kathiresan

**Author notes:** Corresponding authors: Yu-Xin Xu, Center for Genomic Medicine, Massachusetts General Hospital, Simches 5.500, 185 Cambridge St., Boston, MA 02114, USA. Sekar Kathiresan, Center for Genomic Medicine and Cardiovascular Research Center, Massachusetts General Hospital, Simches 5.252, 185 Cambridge St., Boston, MA 02114, USA.

## Abstract

Human genetics studies have uncovered genetic variants that can be used to guide biological research and prioritize molecular targets for therapeutic intervention for complex diseases and metabolic conditions. We have identified a missense variant (P746S) in *EDEM3* associated with lower blood triglyceride (TG) levels in >300,000 individuals. Functional analyses in cell and mouse models show that EDEM3 deficiency strongly increased the uptake of very low-density lipoprotein and thereby reduced the plasma TG level, as a result of up-regulated expression of LRP1 receptor. We demonstrate that EDEM3 deletion up-regulated the pathways for RNA and ER protein processing and transport, and consequently increased the cell surface mannose-containing glycoproteins, including LRP1. Metabolomics analyses reveal a cellular TG accumulation under EDEM3 deficiency, a profile consistent with individuals with carrying *EDEM3* P746S. Our study identifies EDEM3 as a regulator of blood TG, and targeted inhibition of EDEM3 may provide a complementary approach for lowering elevated blood TG concentrations.

## Introduction

High blood triglyceride (TG) level is one of the leading risk factors for coronary artery diseases (CADs) (Bauer et al., 2016; Johansen et al., 2011), along with low-density lipoprotein cholesterol (LDL-C). Human genetic studies have suggested that high TG levels causally affect the risk of CADs independent of LDL-C and high-density lipoprotein cholesterol (HDL-C) (Do et al., 2013). Additionally, diabetes mellitus and metabolic syndrome are two major risk factors for CADs (Malik et al., 2004). High blood TG level linked to insulin resistance is a phenotype common to these metabolic diseases. Thus, lowering blood TG level could further reduce the incidence of CADs as well as type 2 diabetes and metabolic syndrome, especially in high risk individuals. Apolipoprotein (Apo) C3 and Angiopoietin-like 3 (ANGPTL3) are two molecular targets currently under intensive studies for lowering blood TG (Musunuru et al., 2010; Ramms and Gordts, 2018; Xu et al., 2018). The two targets share the common mechanism of inhibition of lipoprotein lipase (LPL) (Larsson et al., 2013; Lee et al., 2009; Ono et al., 2003). Finding other targets independent of LPL could further lower blood TG levels.

Endoplasmic reticulum (ER)-degradation alpha-mannosidase like protein 3 (EDEM3) plays a crucial role in the ER-associated degradation (ERAD) pathway (Eriksson et al., 2004; Hirao et al., 2006; Olivari and Molinari, 2007). Nascent polypeptides emerged in the ER undergo several quality control processes assisted by ER chaperones and folding factors that ensure proper folding and maturation. Misfolded or unassembled proteins are recognized and subject to degradation (Bernasconi and Molinari, 2011; Hebert et al., 2005; Trombetta and Parodi, 2003). The ER-resident EDEM family of proteins is responsible for identifying misfolded proteins by recognizing the N-linked glycans and signaling them for degradation (Bernasconi and Molinari, 2011). EDEM3 is the only EDEM protein that contains α1,2-mannosidase activity, which appears to play a role in its activity to accelerate the misfolded protein degradation. It remains unclear how EDEM3 recognizes the misfolded protein, but previous findings have indicated that EDEM3 does not necessarily target only misfolded glycoproteins, as it has been shown to stimulate mannose trimming amongst the total glycoproteins (Hirao et al., 2006). The biological significance of EDEM3 is still unknown.

We provide evidence that EDEM3 may be another TG-lowering target that represents a distinct mechanism in modulating TG metabolism. We identified a rare variant (P746S) in the coding region of EDEM3 using human genetic studies that is specifically associated with lower blood TG. Deletion of EDEM3 gene in two types of human hepatoma HepG2 and Huh7 cells strongly increased the uptake of VLDL as a result of increased LRP1 expression. Consistent with this observation, knock down of hepatic EDEM3 with CRISPR/Cas9 decreased the plasma TG level in mice. Interestingly, we found that EDEM3 gene deletion increased the cell surface mannose presence without increasing the sensitivity of the cells to ER stress, likely due to the compensation by increased RNA processing and ER protein transport, and repressed cellular metabolic processes. Importantly, the cellular lipid metabolite profile under EDEM3 gene deletion largely explains the plasma lipid metabolite results from the individuals carrying the EDEM3 missense variant, P746S. Thus, our findings show that by regulating LRP1 expression, EDEM3 is able to modulate the plasma TG level. Inhibition of EDEM3 expression could offer an alternative path to lower TG levels.

## Results

### A missense mutation in EDEM3 is linked to lower human plasma TG levels

To investigate possible roles of EDEM3 in lipoprotein metabolism, we first took a human genetic approach to identify EDEM3 single nucleotide polymorphisms (SNPs) that might be associated with lipid traits. We analyzed >300,000 individuals of mainly of European ancestry with exome array genotyping available from the Global Lipids Genetics Consortium (GLGC) against plasma lipid levels (Liu et al., 2017). Among 7 EDEM3 SNPs examined, only one, rs7844298 (P746S), shows statistically significant relations with lower plasma TG level, but not with HDL-C and LDL-C (sTable 1 and Fig. 1A). Comparing this SNP with other validated missense variants shows that its effect on TG level is ∼65% of ANGPTL3 rs77871363 (M259T) and ∼38% of PCSK9 rs28362286 (gain-of-function), both of which exhibit the strong link with TG phenotypes (Fig. 1B). The SNP is located at the coding region of EDEM3 protease associated (PA) domain (Fig. 2C), which is not present in other EDEM members (Olivari and Molinari, 2007). The SNP’s unique domain location and its effect on blood TG concentrations at the population level suggest that EDEM3 may have a novel mechanism in regulating lipoprotein metabolism.

**Fig. 1.**
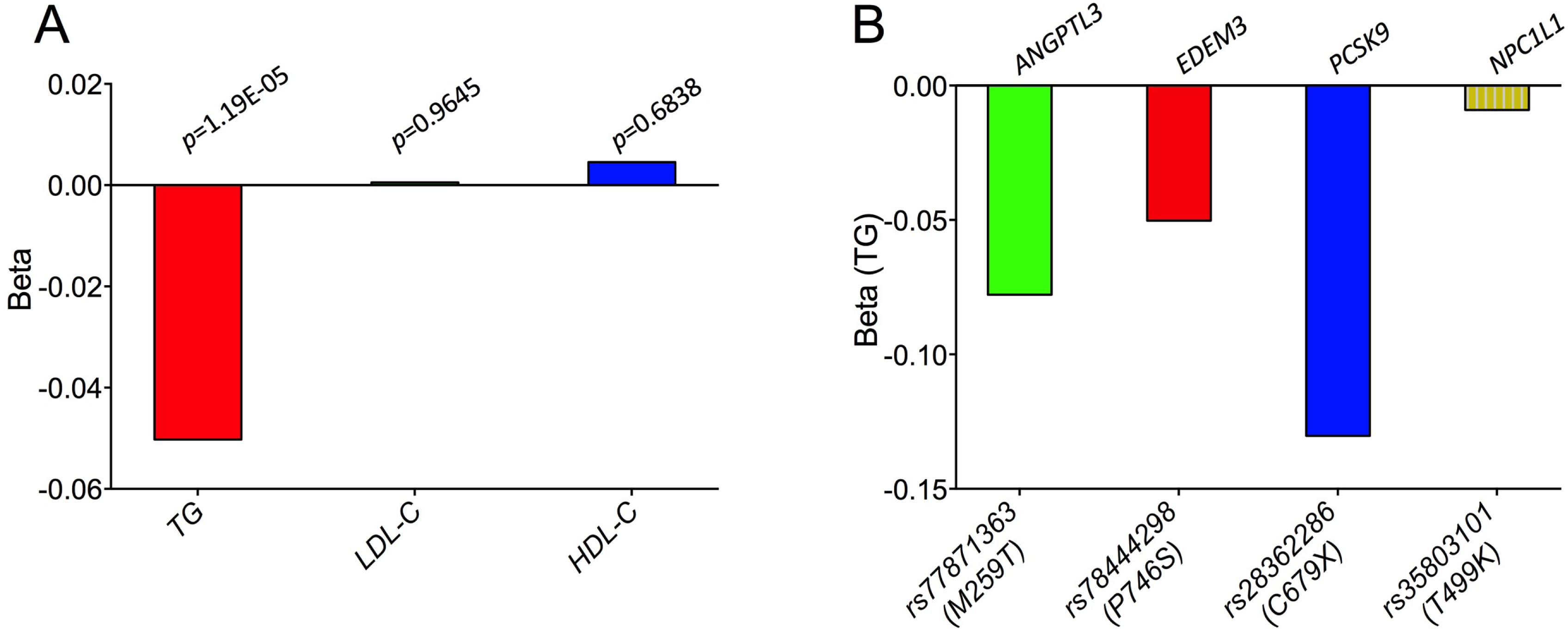
Genetic association study of EDEM3 missense mutations with lipid phenotypes. **A**. Significant association of EDEM3 SNP rs78444298 (P746S) with blood TG (5% decrease; p=1.2X10^-5^), but not with LDL-C and HDL-C (p>0.05). **B**. Comparison of the effect of *EDEM3* rs78444298 with other missense mutations of TG genes including *ANGPTL3* rs77871363, *PCSK9* rs28362286 (gain-of-function) and *NCP1L1* rs35803101 (negative control) on plasma TG. Beta is displayed in standard deviation units of log(TG).

**Fig. 2.**
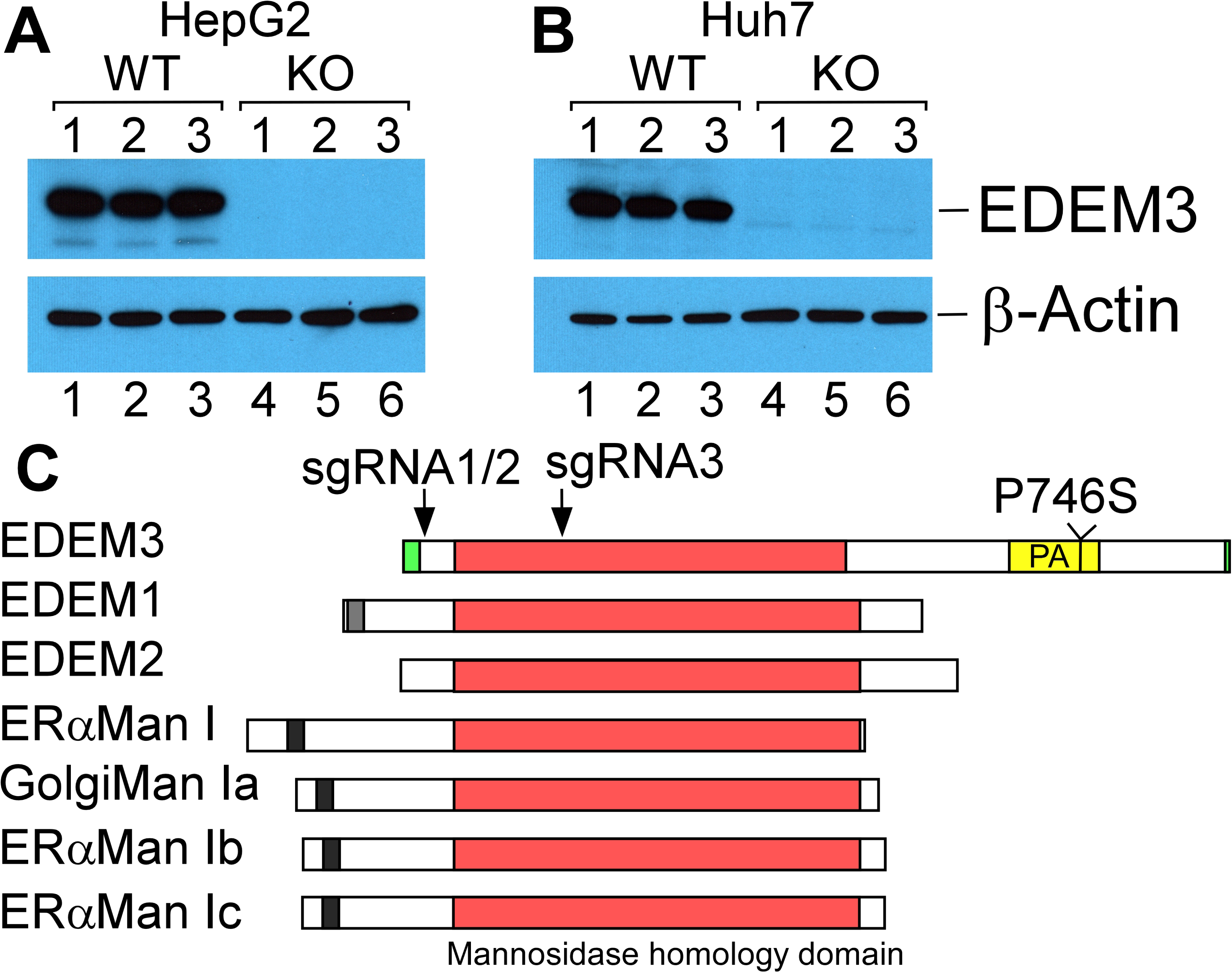
Confirmation of EDEM3 gene deletion in HepG2 and Huh7 cells. **A**. Western analysis of EDEM3 expression. Extracts from EDEM3 WT and KO HepG2 and Huh7 cells were examined with Western blotting and probed with anti-EDEM3 and -β-Actin antibodies. **B**. Domain similarity of Class I a1,2-mannosidase family members. The conserved mannosidase homology domain and the unique protease-associated (PA) domain of EDEM3 are highlighted. The positions of sgRNAs and the human EDEM3 P746S mutation are indicated.

### Deletion of EDEM3 gene strongly increased VLDL, but not LDL, uptake

It is likely that the lower plasma TG level observed in human genetic studies is from the damaging effect of the mutation on the cellular activity of EDEM3. Therefore, to obtain the mechanistic insight of EDEM3 function in lipid metabolism, we tried to delete the gene in human hepatoma cells for functional assays. We used CRISPR/cas9 genome editing system with three sgRNAs that target the exon region of EDEM3 gene (Fig. 2B) along with scramble control sgRNAs. After screening several hundreds of clones, we obtained EDEM3 KO Huh7 and HepG2 cells as well as the scramble controls (WT). Western analysis confirms the deletion of EDEM3 gene specifically in three HepG2 and Huh7 KO clones (Fig. 2A and 2B).

The lower TG phenotypes caused by the EDEM3 mutation could be due to enhanced uptake of TG-rich lipoproteins (TGLs), reduced secretion of VLDL, or both. We first tested whether the EDEM3 gene deletion might affect the VLDL uptake. The KO and control Huh7 and HepG2 clones were incubated with human Dil-VLDL. We found that EDEM3 deficiency strongly increased the VLDL uptake in both Huh7 (by 1.6 times) and HepG2 (by ∼5.2 times) cells (Fig. 3). Dramatic increases were observed in some HepG2 and Huh7 KO clones (by ∼2.6 and ∼10 times, respectively) (Fig. 3A and 3B, bottoms). Similar assays with human Dil-LDL were performed but no significant changes were observed (sFig. 1). The data suggest that deletion of EDEM3 gene increases VLDL, but not LDL, uptake.

**Fig. 3.**
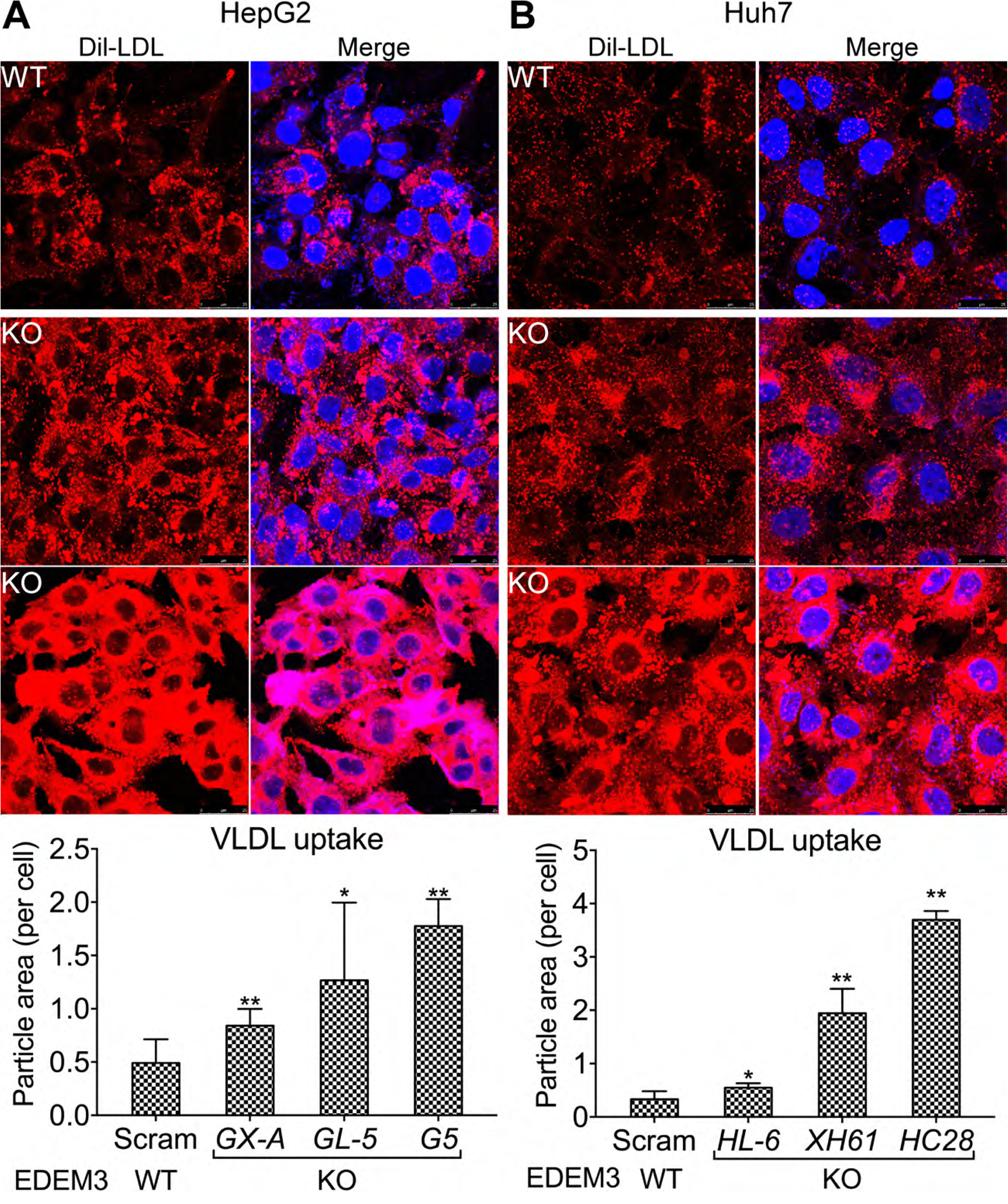
Deletion of *EDEM3* gene strongly enhanced Dil-VLDL uptake. **A**. Three individual EDEM3 scramble control (WT) and KO HepG2 (A) and Huh7 (B) clones were incubated with Dil-VLDL. Tops, Representative images of the uptake assays from *EDEM3* KO and control cells. Bottoms, Quantification of the uptake assays from three clones of the control cells (∼200 cells) and KO cells (∼500 cells). Cellular Dil-VLDL particle areas were quantified using ImageJ. *p* values (t-test) were calculated by comparing the KO cells with the control cells. Bar, 25 μm.

### Deletion of EDEM3 gene did not affect VLDL secretion

We next investigated whether EDEM3 gene deletion might also affect the VLDL or nascent ApoB secretion. To this end, we carried out time course experiments to monitor nascent ApoB-100 secretion and its accumulation in culture media. Three clones of EDEM3 KO and control HepG2 and Huh7 cells were washed and incubated with fresh media. At various time points, fractions of the media were analyzed with ApoB ELISA. The ApoB in the media reflects the dynamics of ApoB secretion from and uptake into the cells. However, at the earliest time points (i.e., 20 and 40 min.), the ApoB in the media most likely represents the nascent ApoB secreted from the cells. We found no significant difference (p>0.05) in the ApoB levels between the KO and control clones in both HepG2 and Huh7 cells at 20 and 40 min time points (Fig. 4A and 4B). We then monitored the ApoB concentrations from Huh7 cells for longer period. After 40 min, the ApoB level in the media of Huh7 cells decreased at various time points, and became significantly decreased (p<0.05) for all the KO clones at 8 hrs (Fig. 4C). Because EDEM3 gene deletion did not affect the ApoB secretion, the increased uptake is most likely responsible for all the decreases of the ApoB in the media. These data are consistent with those from the Dil-VLDL uptake assays, indicating that enhanced VLDL uptake might play a major role in reducing the TG level under EDEM3 deficiency.

**Fig. 4.**
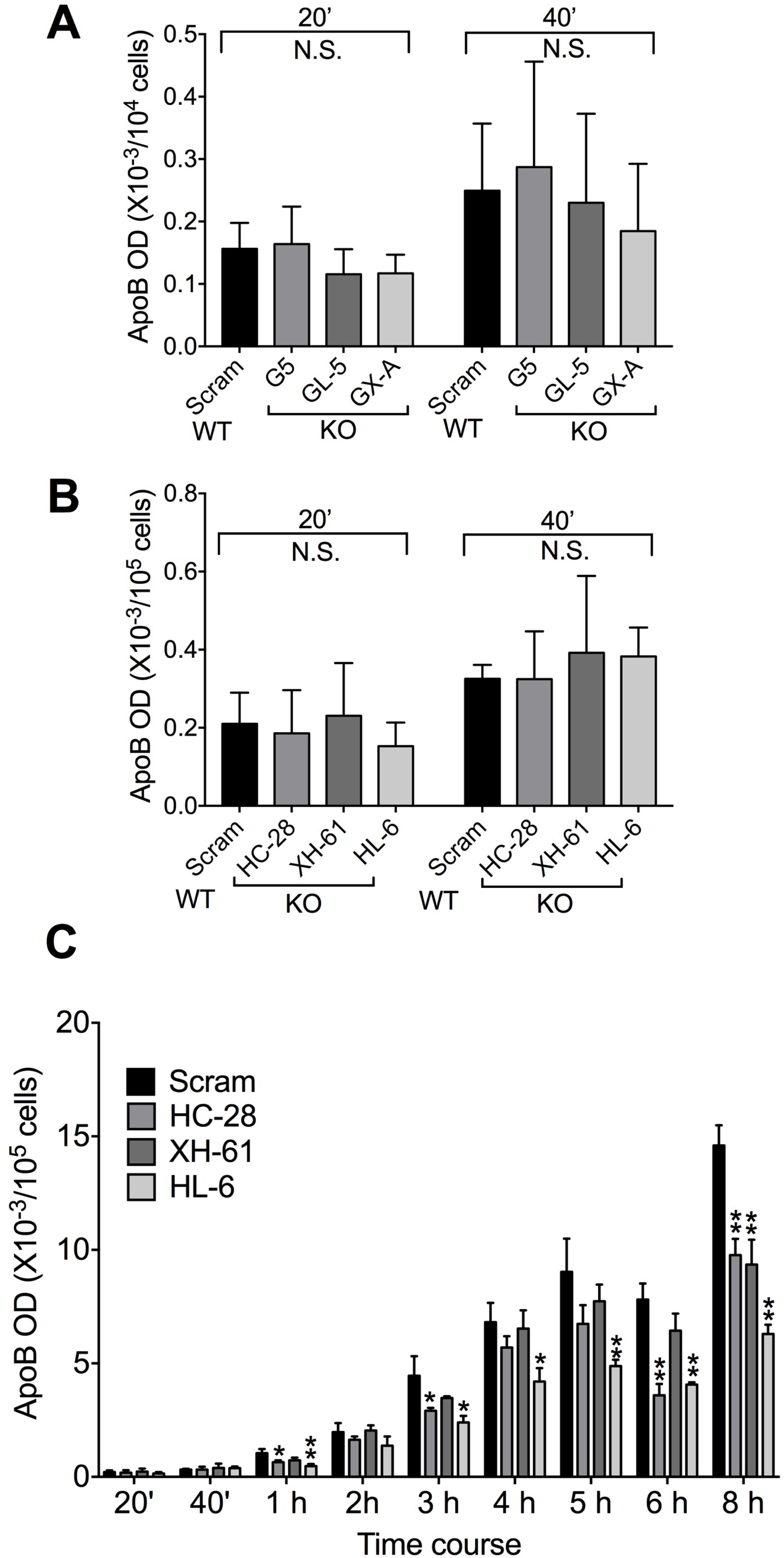
Deletion of *EDEM3* gene reduced ApoB-100 accumulation in culture media without interfering nascent ApoB secretion. Three individual EDEM3 scramble control (WT) and KO HepG2 (**A**) and Huh7 (**B**) clones were used for time course experiments to monitor the ApoB presence in culture media between 20 and 40 mins using ApoB ELISA kit. **C**. Complete time course experiment for monitoring culture media ApoB concentrations from Huh7 cells between 20 min and 8 hrs. *p* values (t-test) were calculated by comparing the KO cells with the control cells.

### Deletion of EDEM3 gene increased LRP1 expression

Hepatic LRP1 plays a central role in clearance of TGLs such as chylomicron and VLDL remnants (Lillis et al., 2008). It is possible that EDEM3 modulates the plasma TG level via regulating the LRP1 expression in the livers. To test this hypothesis, we first used indirect immunofluorescence to examine the cell surface LRP1 expression. The EDEM3 WT and KO HepG2 and Huh7 clones were stained with anti-LRP1 and -ApoB antibodies. The results presented in Fig. 5 show that deletion of EDEM3 gene substantially increased the cell surface LRP1 expression both in Huh7 and HepG2 cells. In the WT control cells, the cell surface LRP1 is barely detectable under the conditions we tested. But in the KO cells, the LRP1 signal is strongly increased (by ∼3.5 times and ∼1.7 times increases in Huh7 and HepG2 cells, respectively). The accumulated LRP1 on the cell surface can be clearly seen at the cell peripheries in the flatter KO Huh7 cells (Fig. 5A).

**Fig. 5.**
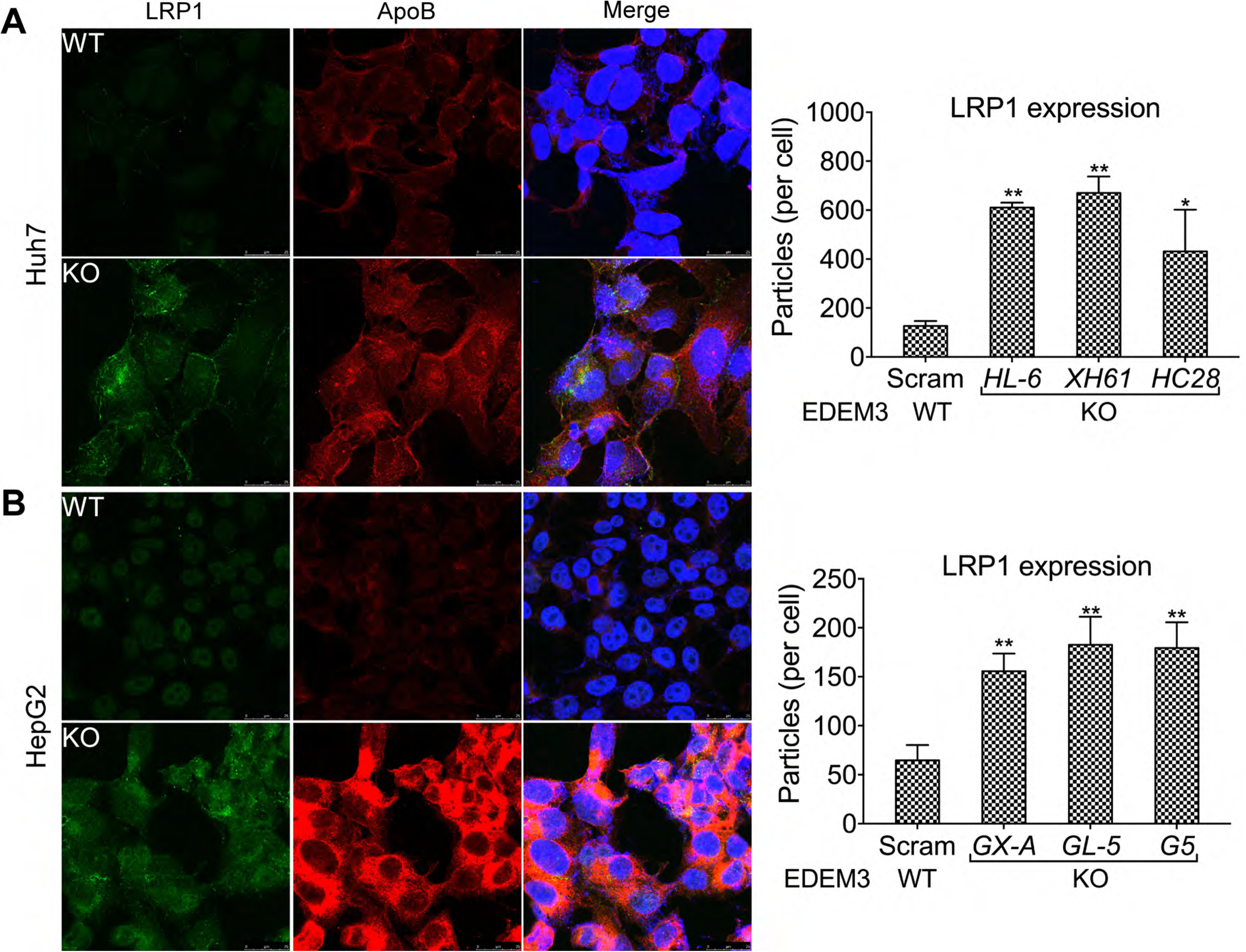
Deletion of *EDEM3* gene increased LRP1 expression. (**A**) Three individual EDEM3 scramble control (WT) and KO Huh7 (**A**) and HepG2 (**B**) clones were stained with anti-LRP1 and -ApoB antibodies, and analyzed with confocal microscope (Left). Images were quantified with ImageJ from 100-200 cells for each clone (Right). *p* values (t-test) were calculated by comparing the results from the KO cells with those from the control cells. Bars, 25 μm.

We then examined the LRP1 expression with Western analysis. Consistent with the above immunofluorescence results (Fig. 5), it shows that the LRP1 protein level is indeed greatly increased in the EDEM3 KO HepG2 and Huh7 cells (Fig. 6A and 6B). Interestingly, we observed that the LRP1 from the KO cells has slower mobility, which is more obvious from the HepG2 KO cells (Fig. 6A, lanes 4-6), indicating that the LRP1 may possess post-translational modification changes related to the activity of EDEM3 in the ER. Because of mannose hydrolase activity of EDEM3, we expected that the LRP1 in the KO cells might retain mannose/glucose, which are supposed to be removed in the ER (Olivari and Molinari, 2007). To test this, the EDEM3 WT and KO HepG2 clones were stained with FITC-labeled Concanavalin A (ConA) (recognizing mannose) and analyzed with flow cytometry. The results show that the KO cells indeed contained stronger signal than the WT cells (Fig. 6C). Thus, the data shown here suggest that deletion of EDEM3 gene increase the cell surface LRP1 expression, which is most likely related to its post-translational processing in the ER.

**Fig. 6.**
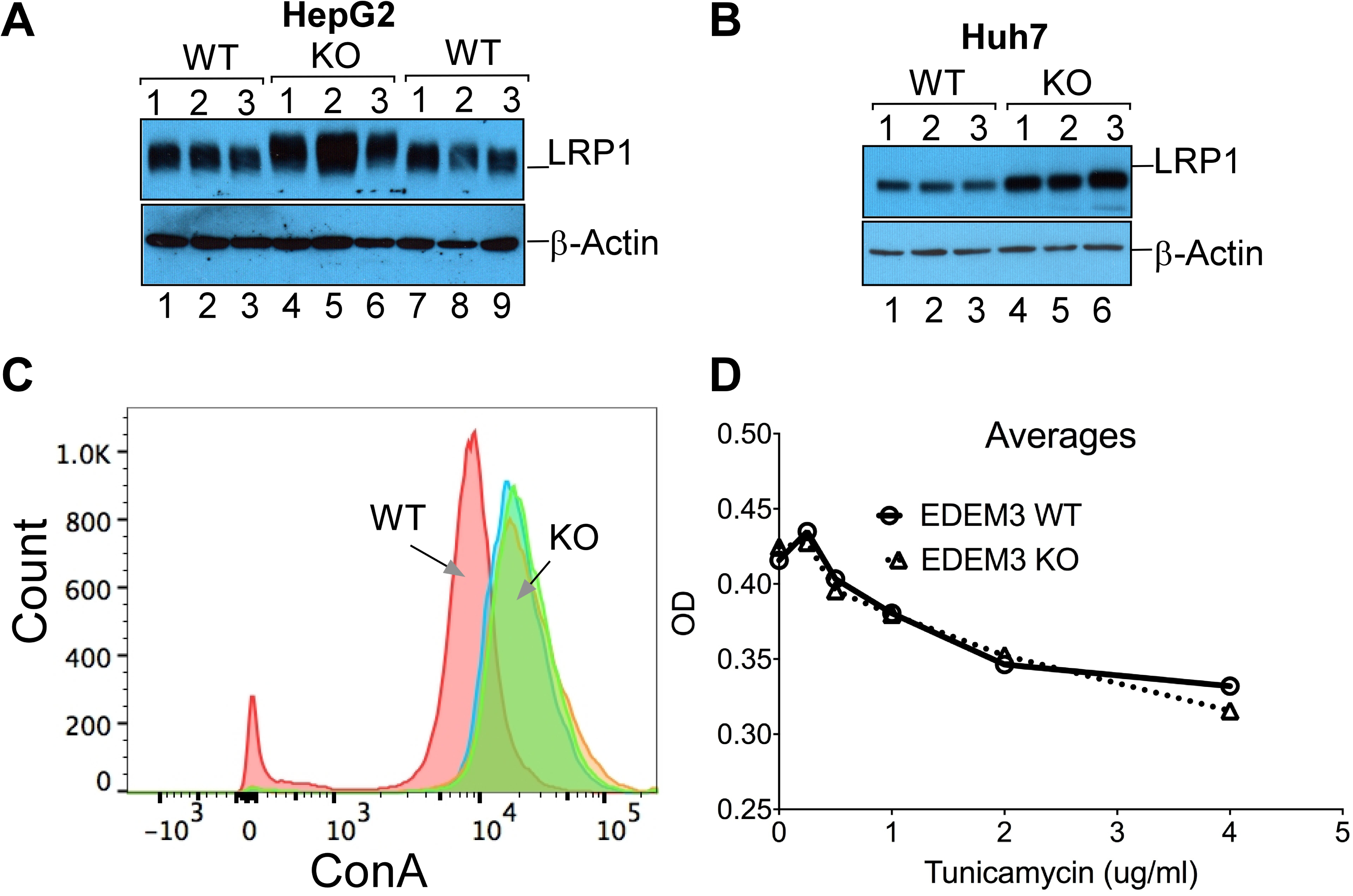
Deletion of *EDEM3* gene increased LRP1 expression as well as cell surface mannose-containing proteins. (**A** and **B**). Extracts from three individual EDEM3 scramble control (WT) and KO Huh7 (**A**) and HepG2 (**B**) clones were analyzed with Western blotting and probed with anti-LRP1 and -β-actin antibodies. **C**. The WT and KO HepG2 clones as in A were stained with FITC-ConA and fixed with 1% paramaldehyde. The cells were then analyzed with flow cytometry. **D**. The WT and KO HepG2 clones as in A were incubated in media without or with increasing amount of tunicamycin as indicated. Viability of the cells were measured using alamarBlue reagent and absorbance was monitored at 570 nm.

### Complement of EDEM3 deletion with WT, but not P746S mutant, EDEM3 reduced LRP1 expression and VLDL uptake

To obtain further functional evidence of EDEM3 in regulating LRP1 expression and VLDL uptake, we performed complementary analysis of EDEM3 deletion with WT and P746S mutant EDEM3. The *EDEM3* KO HepG2 and Huh7 cells were infected with control lentivirus alone or the lentiviruses harboring the cDNAs of flag-tagged WT and P746S mutant *EDEM3*. Western analysis (Fig. 7A, middle) confirmed the expression of the WT (Fig. 7A, middle, lanes 4-6) and mutant (7-9) *EDEM3*. The expression of WT EDEM3 significantly reduced the level of LRP1 (by ∼11%). Whereas, the mutant *EDEM3* even increased the LRP1 expression by ∼74%. The LRP1 from the KO cells with WT EDEM3 complement had slightly faster mobility as compared with those from the KO cells and the cells with the mutant EDEM3, indicating of modification changes. Immunofluorescence analysis confirmed the LRP1 expression changes in the KO HepG2 cells and the cells with the WT and mutant EDEM3. The increased LRP1 expression correlated with the elevated cellular ApoB signal (sFig. 2). Similar results were also obtained from the Huh7 KO with the WT and mutant EDEM3 (sFig. 3).

**Fig. 7.**
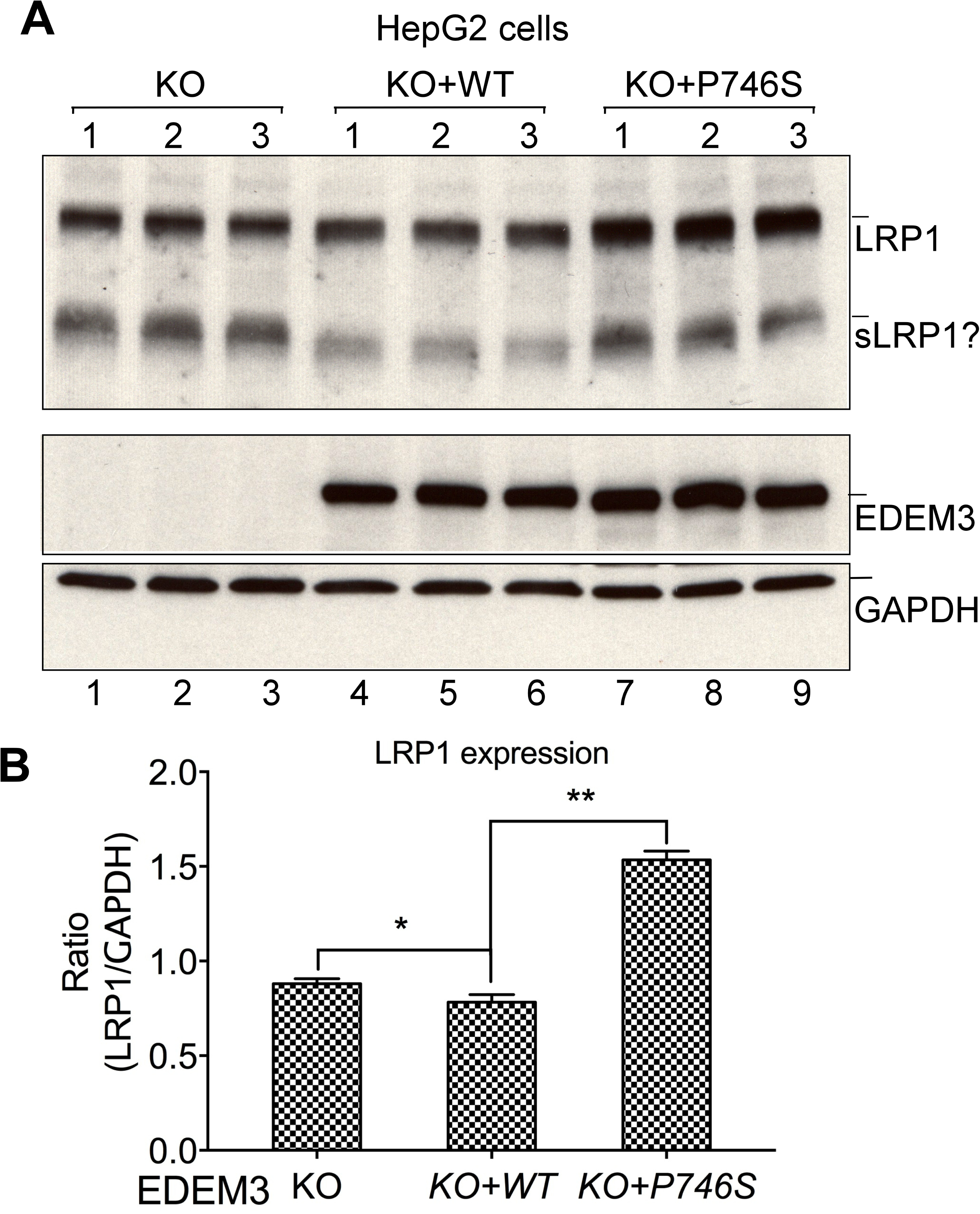
Complementary analysis of *EDEM3* gene deletion with WT and P746S mutant EDEM3. **A.** EDEM3 KO HepG2 cells were infected with control lentivirus or the same virus containing WT and P746S EDEM3 cDNA. Extracts from the complementary or control cells were analyzed with Western blotting and probed with the antibodies against LRP1, EDEM3 and GAPDH (from top to bottom). **B**). Normalized quantification of LRP1 expression with GAPDH as in A. *p* values (t-test) were calculated by comparing the KO+WT cells with the KO or KO+P746S cells. ANOVA *p* value<0.001. Bar, 25 μm.

The above cells were then used for VLDL uptake assays. The complementary expression of WT EDEM3 drastically reduced the VLDL uptake (by ∼62%) as compared with those of the KO HepG2 cells. In contrast, the expression of the mutant EDEM3 in the KO cells increased the VLDL uptake (by nearly two times) as compared with the WT complement (sFig. 4). Again, similar results were obtained from Huh7 KO cells and the cells with complementary WT (decreased by ∼28%) and mutant (increased by ∼57%) EDEM3 expressions (sFig. 5). The VLDL uptakes in both cell types are correlated to the expression levels of LRP1. Therefore, the above results highlight the crucial role of EDEM3 in regulating LRP1 post-translational processing, and deletion of EDEM3 gene or the loss-of-function mutation facilitate the processing and transport of LRP1 to the cell surface.

### Hepatic EDEM3 KD in mice increased LRP1 expression and essentially reduced plasma TG level

To obtain in vivo evidence for the regulation of EDEM3 in LRP1 expression, we knocked down EDEM3 expression in mouse livers using CRISPR/Cas9 genome editing system (Ran et al., 2015). We selected one sgRNA from in vitro screening a pool of sgRNAs that target EDEM3 exon 1 (see Materials and Methods). The sgRNA was cloned into a single vector adeno-associated virus (AAV) -Cas9 system containing *Staphylococcus aureus* Cas9 (SaCas9). The viral constructs containing the EDEM3 sgRNA and a non-specific scramble sgRNA (control) were then used for AAV production. The AAV preparations were injected in WT female mice followed by high fat diet.

Western analysis shows that the CRISPR-based KD reduced the hepatic EDEM3 expression (by ∼50%) (Fig. 8A, top and 8B, left). And the EDEM3 KD essentially reduced the plasma TG level on day 18 and 27 (by ∼43% and ∼29%, respectively) (Fig. 8C and 8D). The higher plasma TG observed at days 18 and 27 in the control mice was likely induced by the high fat diet. The TG reduction suggests that inhibition of EDEM3 might be able to decrease the diet-induced high TG level. Consistent with the above results, EDEM3 KD significantly increased (by ∼50%) the expression of hepatic LRP1 (Fig. 8A, middle and 8B, right). Similarly, we also observed some detectable mobility changes of the small subunit of LRP1 as more molecules of LRP1 from the EDEM3 KD mice were shifted to slightly higher molecular weight (the mobility change is more obvious in the gel with less protein loading, Fig. 8A, bottom), indicating the modification change of LRP1 indeed occurred in vivo. Thus, we provide both in vitro and in vivo evidences that EDEM3 inhibition increases the expression of LRP1, which in turn increases the TG-rich lipoprotein uptake and reduces the plasma TG level.

**Fig. 8.**
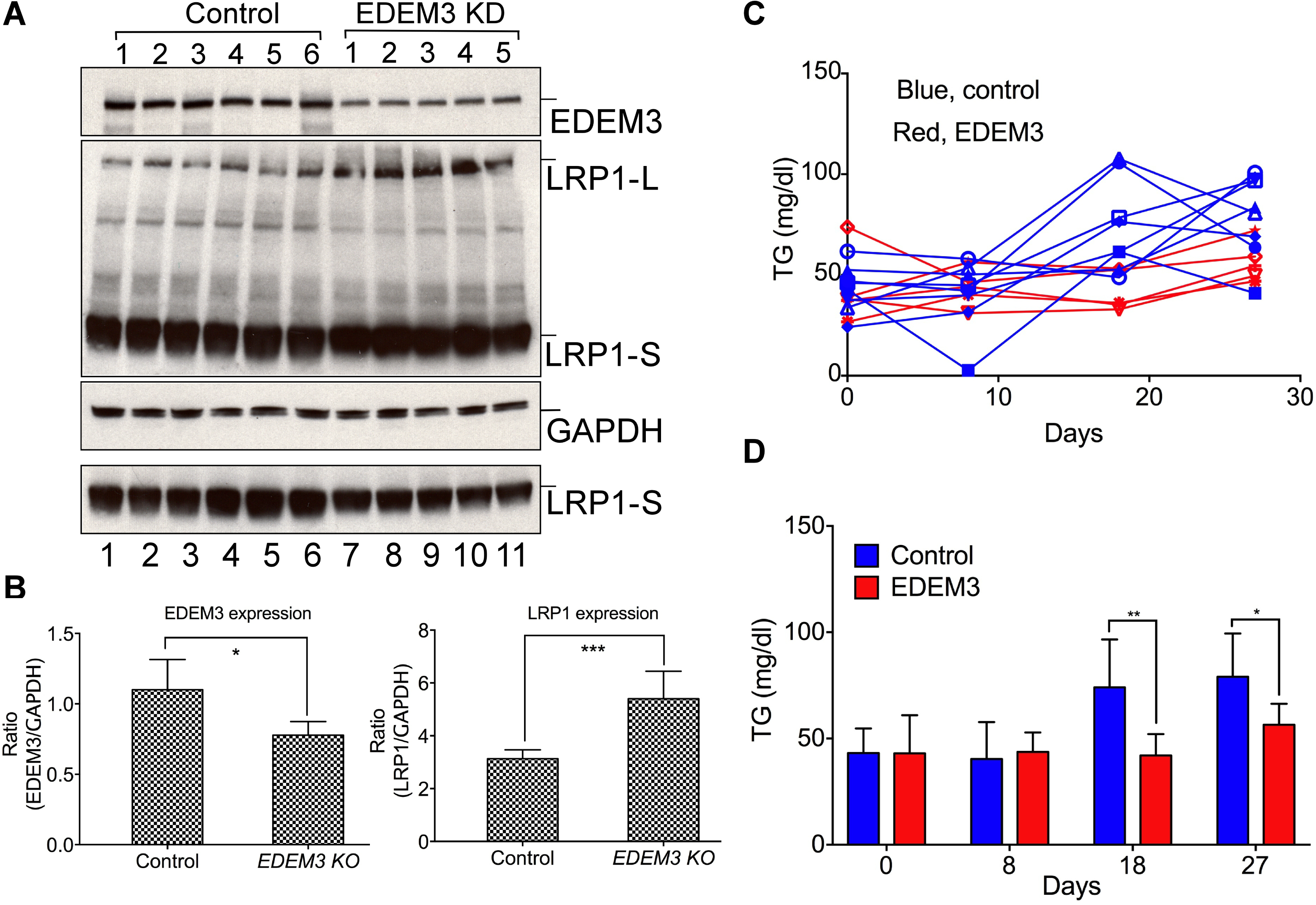
Hepatic *EDEM3* KD increased LRP1 expression and reduced the plasma TG level in mice. **A.** Liver extracts from the female mice injected with the control AVV or the AAV for EDEM3 CRISPR KD were analyzed with Western blotting and probed with anti-EDEM3, -LRP1 and –GAPDH antibodies (from top to bottom). Note: Bottom panel, the amount of the extracts for lanes 7-11 was 1/3 of those for lanes 1-6. **B**). Quantification of EDEM3 and LRP1 expression normalized with GAPDH as in A. **C** and **D**. Measurement of the plasma TG levels from the mice injected with the AAVs as in A. **C**. Individual mouse plasma TG levels; **D**. Average plasma TG of the control and EDEM3 KD groups. *p* values (t-test) were calculated by comparing the KD mice with the corresponding control mice.

### EDEM3 deletion increased the expression of the genes required for RNA and protein processing and transport

EDEM3 functions in protein quality control in the ER (Olivari and Molinari, 2007). Deletion of EDEM3 gene might affect the sensitivity of cells to ER stress. To test this, we treated EDEM3 KO and control HepG2 cells with tunicamycin, a reagent known to induce ER stress (Graham et al., 2016). The results (Fig. 6D) show that the treatment did not induce the cell viability changes between the KO and control clones. We observed slightly more cell deaths only in one clone at the highest concentration of tunicamycin. Thus, the deficiency of EDEM3 might not add too much burden on protein processing in the ER.

It is likely that the KO cells might adapt some gene expression changes that enable them to tolerate the deficiency of EDEM3. To investigate this, we performed RNA-seq analysis on the total RNA from the EDEM3 KO and WT HepG2 clones. Differential gene expression analysis shows that the EDEM3 gene deletion induced down-regulation of expression of a large number of genes (Fig. 9A). The down-regulated gene expression resulted in reduced activities of many cellular metabolic pathways such as PPAR signaling, drug metabolism, and protein digestion and absorption, etc. (Fig. 9B). Importantly, the pathway analyses show that the expression of the genes for RNA and protein processing and transport are up-regulated (Fig. 9C and 9D). Thus, the up- and down-regulated pathways may directly alleviate the burden potentially resulted from the loss of EDEM3 protein in the ER, and facilitate the LRP1 post-translational processing in the ER and transport on the cell surface.

**Fig. 9.**
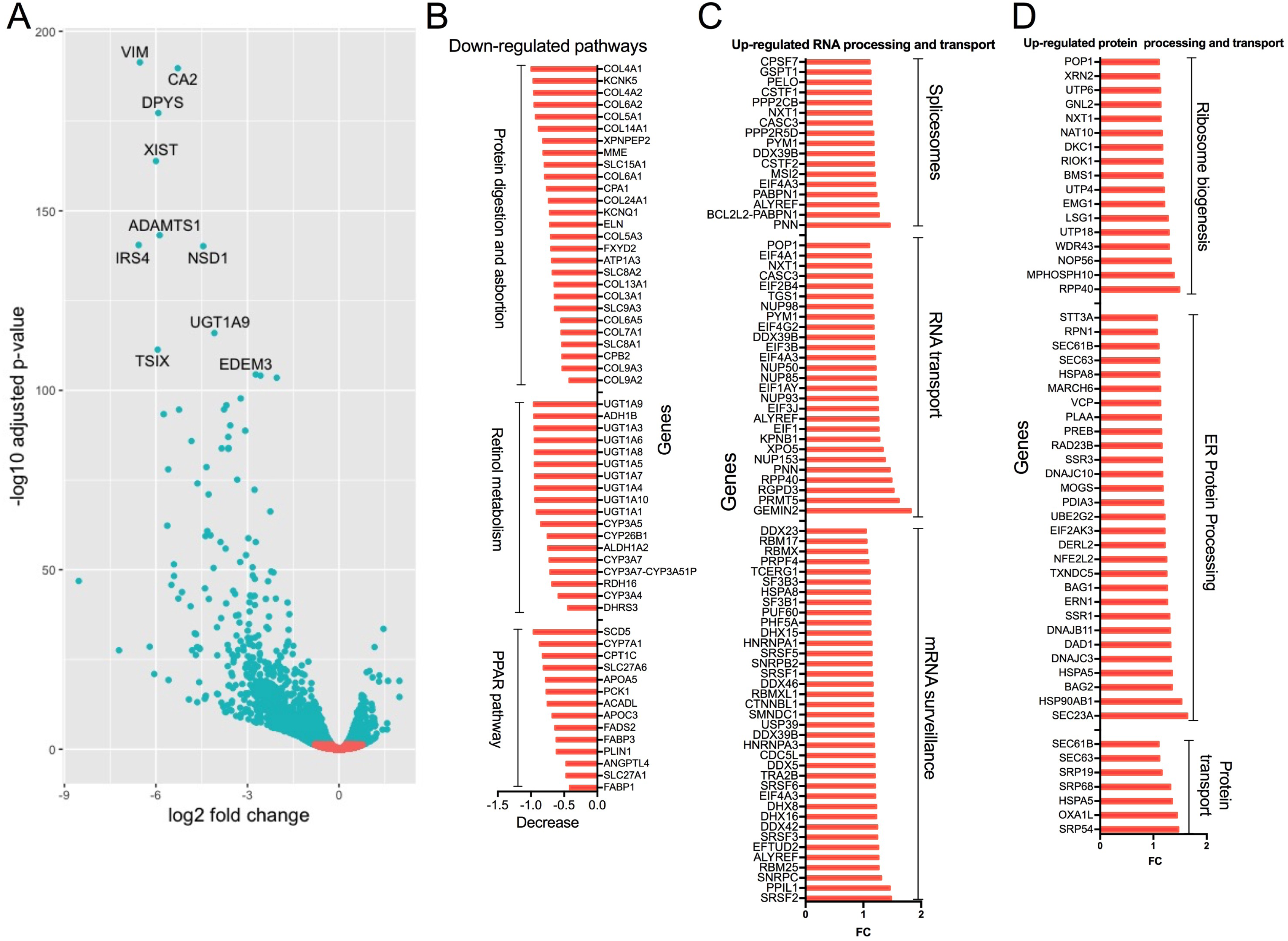
Deletion of *EDEM3* gene in HepG2 cells increased RNA and ER protein processing and transport. Total RNA from three individual EDEM3 scramble control (WT) and KO HepG2 clones was analyzed with RNA-seq for differential gene expression (**A**). The down-regulated (**B**) and up-regulated (**C** and **D**) pathways related to EDEM3 gene deletion were shown.

### Plasma metabolite profile of rs78444298 (EDEM3 P746S) corresponds to the cellular metabolite pattern of EDEM3 gene deletion

To investigate the lipid metabolite changes resulting from EDEM3 gene deletion, we used mass spectrometry to analyze the total lipid extracts from the EDEM3 KO and control HepG2 cells as we previously reported (Xu et al., 2018). The results show that the gene deletion induced the accumulation of some DAGs and shorter carbon chain TAGs (Fig. 10A and 10B). For longer carbon chain TAGs, some of them increased, while others decreased, however, if the cellular metabolite abundance is taken into consideration, there is still accumulation of the longer carbon chain TAGs (Fig. 10A). For instance, there are extremely low levels of C58:10 and C58:11 TAGs. These TAGs are much less abundant than those elevated TAGs including C50:3, C52:1, C54:1, C54:2, C56:2 and C56:3. There is no significant change for the most abundant C52:2 TAG.

**Fig. 10.**
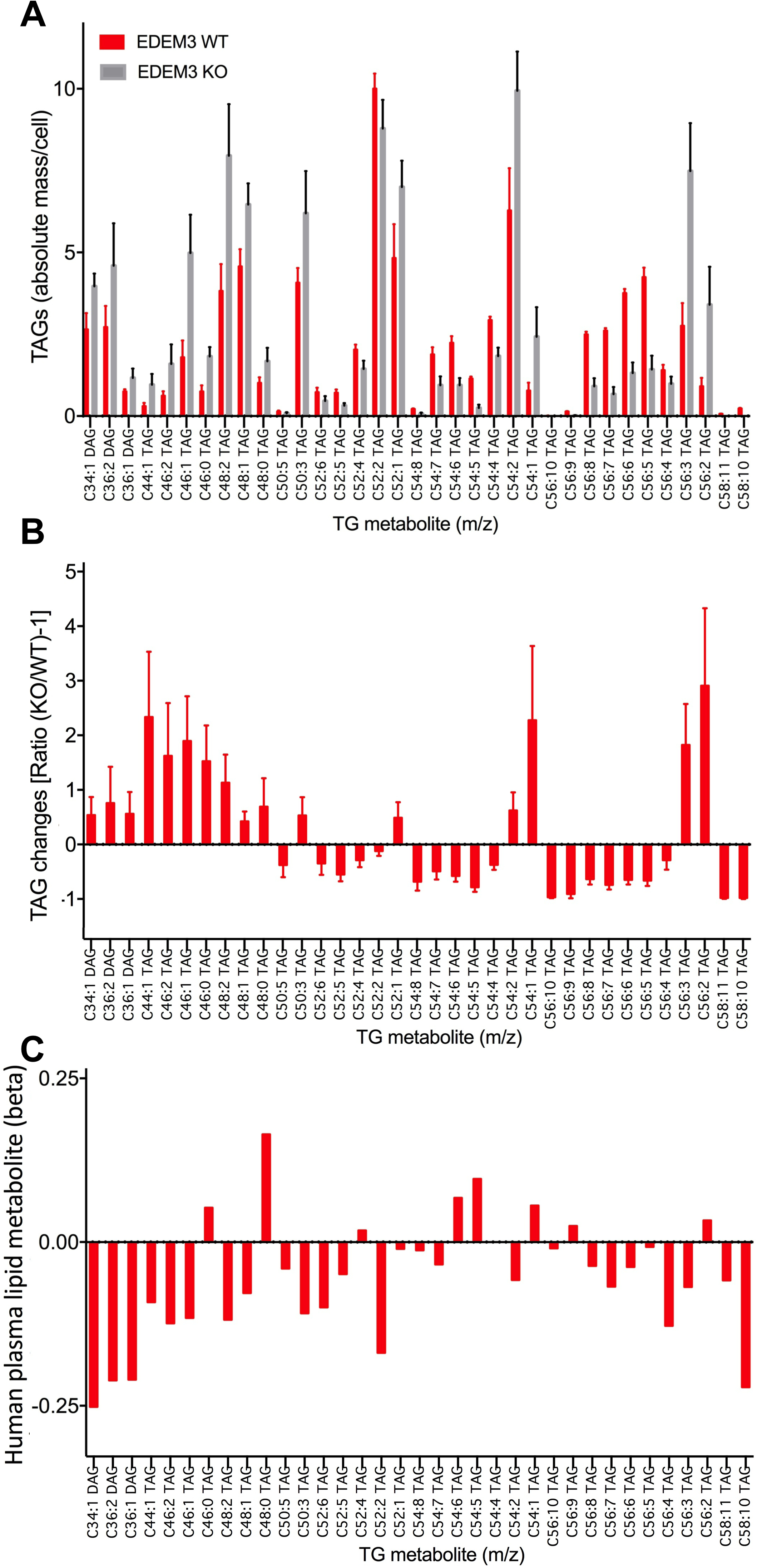
Comparison of the lipid metabolite profiles of rs78444298 (EDEM3 P746S) FHS individuals with those from *EDEM3* gene deletion in HepG2 cells. **A**. Total cellular lipid extracts from three individual EDEM3 scramble control (WT) and KO HepG2 clones was analyzed with mass spectrometry. The metabolite profile was obtained by comparing the data from KO with those from WT cells. **B**. The association plasma lipid metabolite profiles in 1797 individuals from FHS with rs78444298 (P746S) from FHS 1000G GWAS cohorts.

We next examined the plasma lipid metabolite changes by SNP rs78444298 genotype (G>A; 1.5% minor allele frequency) in 1797 individuals from the Framingham Heart Study (FHS) with 1000 genomes imputated genotypes (Rhee et al., 2013). The results demonstrate a similar, but in opposite direction, pattern of DAGs and shorter carbon chain TAGs as compared with those from EDEM3 gene deletion (Fig. 10B and 10C). More decreases were found for the longer carbon chain TAGs. The longer and abundant TAGs (C50:3, C52:1, C54:2, and C56:3), which were accumulated in the EDEM3 KO HepG2 clones, are decreased in increasing number of A alleles carried. Thus, in general the plasma lipid metabolite profile with increasing number of A alleles is correlated with the corresponding pattern change of the EDEM3 gene deletion, in particular for those DAGs and shorter carbon chain TAGs. The decreased DAGs and TAGs might result from the increased uptake by the livers because of elevated LRP1 expression induced by the LoF EDEM3 mutation. However, more in vivo studies are needed to explore the lipid metabolic changes.

## Discussion

We demonstrate that an EDEM3 variant, P746S, is significantly associated with the lower blood TG level using human genetic data. We sought to use a cell-based functional approach by deleting the EDEM3 gene in human hepatoma cells to obtain mechanistic insight for this association. We found that the gene deletion strongly induced the uptake of VLDL as a result of up-regulated LRP1 expression. Hepatic EDEM3 KD in mice further confirmed the results in vivo. RNA-seq analyses reveal that the gene deletion up-regulated the pathways for RNA and ER protein processing, which likely facilitated the LRP1 processing in the ER and transport to the cell surface. Furthermore, our metabolomics data show that the cellular lipid metabolite profile is largely consistent with the counterpart association of the missense variant in the plasma of individuals from the population, suggesting that our functional analyses largely explain the phenotype from the genetic association. Collectively, we identified EDEM3 as a regulator in modulating plasma TGs via its activity in the ER and thus it may represent a potential therapeutic target for TG lowering.

Blood TG level represents the TG content of TGLs in circulation, mainly including the remnants of chylomicron from dietary intake and VLDL secreted from the livers. Blood TG is associated with dietary fat intake, genetic determinants, and physical activity, as well as other factors. There are several drugs available for the treatment of hypertriglyceridemia such as niacin, fibrates and omega-3-fatty acids (Cooper et al., 2015; Group et al., 2018; Pawlak et al., 2015), but none of them is specific. Finding targets for lowering TG and further using them for drug development have been lagged behind the drug development for LDL-C lowering. Recent human genetic studies have identified ApoC3 and ANGPTL3 as two important TG targets (Jorgensen et al., 2014; Musunuru et al., 2010; TG et al., 2014). Inhibition of ApoC3 and ANGPTL3 showed strong effects in lowering the blood TG (Gaudet et al., 2014; Gaudet et al., 2017; Xu et al., 2018). But the two targets share a similar mechanism in modulating the blood TG, i.e. the inhibition of LPL. As mentioned earlier, the three top LDL-C lowering drugs, statin, ezetimibe, and the PCSK9 inhibitor, cover the hepatic cholesterol synthesis, intestinal cholesterol absorption, and LDL uptake, respectively (Ajufo and Rader, 2016). Combination use of these drugs will provide great strength for treatment of complex hyperlipidemia, which can not be controlled by a single therapeutic agent. Thus, the finding of EDEM3 and its role in modulating blood TG via regulating LRP1 expression would complement the current TG-lowering therapeutic efforts.

EDEM3 modulates the plasma TG through regulating LRP1 expression that is consistent with the role of LRP1 in regulating TG metabolism (Lillis et al., 2008). In contrast to the LDLR, which binds to ApoB-100 as well as ApoE, LRP1 mediates the hepatic uptake of the TG-rich lipoproteins mainly through its binding with ApoE, which is present in both chylomicrons and VLDL (Beisiegel, 1998; Mahley and Huang, 2007). It was observed that genetic defects of LDLR both in humans and mice cause hypercholesterolemia by reducing the uptake of cholesterol-rich LDL, but the clearance of TG-rich lipoproteins is not affected (Kita et al., 1982; Rubinsztein et al., 1990). High plasma TG was only observed when the LRP1 expression is inactivated under the LDLR deficiency (Rohlmann et al., 1998; Willnow et al., 1995). Though the mechanisms for LRP1 and LDLR in regulating TG metabolism remain to be determined (Rohlmann et al., 1998), the existing data suggest that LRP1 plays a major role in the clearance of TGLs.

EDEM3 likely regulates the LRP1 expression in a post-transcriptional manner since our RNA-seq analysis did not show significant differences in LRP1 mRNA transcript levels between the EDEM3 WT and KO cells (Fig. 9). It was previously shown that all the LDLR family members including LRP1 possess N-linked glycosylation (May et al., 2003). The mannose-containing N-linked glycans are co-translationally added to the nascent receptor polypeptides, which serves as signals recognized by the ER-resident chaperones and folding factors to ensure proper maturation. The unrecoverable misfolded polypeptides undergo mannose-trimming by the GH47 family mannosidases including EDEM1, 2 and 3, and are then eventually translocated from the lumen of ER into cytoplasm for degradation. EDEM proteins play important roles in maintaining protein folding efficiency through facilitating the extraction of misfolded glycoproteins and accelerating the onset of their degradation. It is intriguing that the EDEM3 deficiency increases the cell surface LRP1 expression. EDEM3 deficiency may accelerate the processing and transport of LRP1 as well as other glycoproteins from the ER. The acceleration may not be related to the quality control of misfolded LRP1 because EDEM3 was demonstrated to carry out the mannose-trimming not only for the misfolded glycoproteins, but for the total glycoproteins as well (Hirao et al., 2006). The other possibility for the increased expression is that the mannose-containing N-linked glycans may increase the stability of LRP1 because glycosylation was shown to promote the stability of LDLR (Pedersen et al., 2014). Further studies are needed to validate these hypotheses.

Our studies indicate that the deletion of EDEM3 gene did not alter the cellular response to the induced ER stress in vitro (Fig. 6D), suggesting that under normal growth condition, there might not be many misfolded proteins accumulated in the ER of the KO cells as compared with the WT cells. This may reflect the change of gene expression resulted from the EDEM3 gene deletion. We observed that the expression of the genes responsible for cellular metabolic pathways was down-regulated, whereas the expression of the genes involved in the RNA and ER protein processing and transport was up-regulated. These gene expression changes may be able to compensate any defects in the ER caused by the loss of EDEM3 protein. However, the expression of the downstream genes regulated by unfolded protein response (UPR) pathways is virtually no difference between the KO and WT cells. For instance, no significant changes were found for the expression of the transcription factor ATF6 and its target genes such BiP, protein disulfide isomerase and glucose-regulated protein 94, which are the key ER-resident proteins for protein folding (Bernasconi and Molinari, 2011; Walter and Ron, 2011). Additionally in a typical UPR, the mRNA translation is inhibited in order to reduce the protein loading in the ER under ER stress. In this case the mRNA and protein production was up-regulated in the EDEM3 KO cells. Thus, the gene expression change may not be related to UPR. The increased RNA and protein production may be the response of the KO cells to compensate the reduced capacity in the protein processing due to the EDEM3 deficiency.

Genetic factors contribute to over 20% of the inter-individual variation of human plasma metabolites (Gieger et al., 2008; Rhee et al., 2013). Given the key roles of metabolites as markers and effectors in cardiovascular diseases, it has become increasingly important to interrogate machineries and mechanisms behind the metabolite variations from population-based cohorts via genetic and biological studies. In this study, we leveraged human metabolomics data in over 1,500 individuals. We compared the association of EDEM3 P746S plasma lipid metabolite profile with the corresponding cellular pattern of the EDEM3 KO cells. They are largely correlated, suggesting that the up-regulated LRP1-mediated TG-rich lipoprotein uptake may have a major impact on the human TG phenotype. Although further studies are needed to obtain a broad spectrum of human plasma metabolite data and expand the studies on the functions of EDEM3 in regulating the cellular lipid metabolism, our study provides an example of how human genetic, functional and mechanistic, and human metabolomics studies can be emphasized for a broad biological insight with respect of disease pathogenesis of a gene.

In summary, in this study, by using a population-based genetic approach, we first demonstrated that a missense mutation in the coding region of EDEM3 is significantly associated with lower plasma TG in a large association study. We then performed both in vitro and in vivo functional analyses by either deleting EDEM3 gene in human hepatoma cell lines or hepatic EDEM3 KD in mice. We observed that the inhibition of EDEM3 expression increased LRP1 expression, which in turn induced TG-rich lipoprotein uptake and further reduced the plasma TG level. Finally, our metabolomics analysis indicate that EDEM3 deficiency increased cellular TG accumulation, which is inversely associated with the lipid metabolite profile of the plasma from the individuals carrying the EDEM3 alternate allele, suggesting that the LRP1-mediated TG-rich lipoprotein uptake plays a major role in the TG phenotype. Importantly, EDEM3 inhibition increased the expression of genes involved in the RNA and ER protein processing and transport, without altering the sensitivity of cells to ER stress, indicating that inhibition of EDEM3 expression could be potentially used as therapeutic approach to treat hypertriglyceridemia.

## STAR Methods

### Cell culture, whole cell extracts, antibodies and Western blotting analysis

EDEM3 KO and WT control Huh7 and HepG2 cells were cultured in DMEM medium (Invitrogen) supplemented with 10% fetal bovine serum (FBS). Whole cell extracts were prepared with the lysis buffer containing 150 mM NaCl, 50 mM Tris (pH 7.5), 1% IGPAL-CA-630 (Sigma # I8896) and protease inhibitor cocktail (Roche). Western blotting analysis were performed using precast Mini-PROTEAN TGX SDS-PAGE (4-20%) (Bio-Rad). Blots were probed with anti-EDEM3 (Sigma, E0409) and -β-actin (Santa Cruz, sc-47778) mouse monoclonal antibodies, and anti-LRP1 (Abcam, ab92544) and -GAPDH (Cell Signaling, 3683S) rabbit monoclonal antibodies.

### *EDEM3* gene deletion using CRISPR/Cas9 genome editing

EDEM3 KO cells were generated as previously described. Briefly, three single guide RNA (sgRNA) oligoes, 5’-GCGAGCGCGATGGAGACTAG-3’, 5’-ATGGAGACTAGTGGCGGCGA-3’, and 5’-CCGGCTTTCAACACTACCAG-3’ were designed to target exon 1 and 5 of human *EDEM3* gene and subcloned into lentiGuide-Puro vector. The vectors containing EDEM3 sgRNAs or a scramble control sgRNA (5’-GCACTACCAGAGCTAACTCA-3’) together with accessory plasmids for lentivirus assembly were co-transfected into 293T cells for lentivirus production (Sanjana et al., 2014; Shalem et al., 2014). Packaged viruses were used to transduce the Cas9-expressing HepG2 and Huh7 cells for ∼16 hrs. The transfected cells were treated with puromycin (5 μg/ml) for five days. Subsequently, single cell clones were screened and the EDEM3 KO was confirmed with Western analysis with anti-EDEM3 and anti-β-actin antibodies.

### ApoB-100 secretion and measurement, and VLDL/LDL uptake assays

Time-course ApoB-100 secretion assays were carried out as previously described. Briefly, EDEM3 KO and control HepG2 and Huh7 cells (3 clones) were splitted and next day the cells were washed with serum-free media and incubated with serum-free medium. At various times media was taken for ApoB-100 measurement using ELISA kit (MABTECH) according to the manufacturer’s instructions. The amount of ApoB-100 was normalized with cell numbers.

For VLDL/LDL uptake assays, EDEM3 KO and control HepG2 and Huh7 cells with, and the KO cells with WT/mutant EDEM3 complementation and relevant controls were incubated in serum-free medium (2 hrs) followed by incubation in serum-free medium containing Dil-VLDL and Dil-LDL (Stephan and Yurachek, 1993) (5 μg/ml, Invitrogen) for 1 hr. The cells were then fixed with 4% paraformaldehyde. The cells were then analyzed under confocal microscope.

### Lentivirus preparation and complementation of EDEM3 KO with WT and P746S mutant EDEM3

Human WT EDEM3 cDNA was purchased from Genescript (Clone ID OHu101917). The P746S mutation was generated using Quickchange site-directed mutagenesis (Agilent Technologies, 200519-5) according to the manufacturer’s procedures. The cDNAs were cloned into the expressing vector FU-tetO-Gateway (Addgene, #43914) using Gateway Technology (Invitrogen) according to the manufacturer’s procedures. The constructs without an insert or containing WT and mutant EDEM3 cDNAs were used for lentivirus production at the viral core of Boston Children’s Hospital. Both the WT and mutant cDNA sequences were confirmed by DNA sequencing. The lentivirus particles were then used to incubate with EDEM3 KO HepG2 and Huh7 cells. Single cell clones were screened for the expression of WT and mutant EDEM3 using Western analysis with anti-EDEM3 antibody.

### Adeno-associated virus (AAV) preparation and CRISPR/Cas9 KD of hepatic EDEM3 expression in mice

A cell based in vitro system was used for screening the sgRNAs targeting mouse EDEM3 exon 1. Briefly, mouse EDEM3 sgRNAs were cloned into pX602 plasmid containing Cas9 from *Staphylococcus aureus* (SaCas9). A genomic fragment containing the target sites of EDEM3 sgRNAs were cloned into pCAG-EGXXFP plasmids. The two plasmids were co-transfected into 293T cells. The EGFP fluorescence intensity in the transfected cells was monitored for the cleavage efficiency of the sgRNAs. A total of 25 EDEM3 sgRNAs were screened. The pX602 constructs containing the most efficient sgRNA (5’-CGGCTGCCGGGGCTGTGGGT-3’) or a scramble control sgRNA (5’-TTTTTTGTTTTTTGTTTTTT-3’) were used for AAV production at the Penn Vector Core. The AAV preparations (total droplet digital genome copy titer, 4.716E+12) were injected into WT mice (C57BL/6J, female, 12 weeks) (Jackson Laboratory) followed by TG-rich high fat diet (Envigo, TD.06414). Blood samples were collected prior to the injection and weekly after the injection. The mice were sacrificed in about one month later. The liver samples were collected.

### Cell surface fluorescein-labeled ConA staining and flow cytometry analysis

EDEM3 KO and control HepG2 cells were blocked with BSA in PBS with 5% FBS and 2 mM sodium azide on ice for 20 min. The cells were then pelleted and resuspended in PBS with fluorescein-labeled ConA (10 ug/ml) (Vector laboratories, FL1001) on ice for 30 mins. The cells were pelleted again and resuspended 250 ul PBS. About 2.5 ml cold paraformaldehyde (1%) was added in dropwise to the cell suspension and incubated on ice for 15 min. The cells were washed twice with cold PBS. The cell pellet was resuspended in 250 ul cold PBS and 1 ml cold 70% ethanol was added in dropwise. The suspension was incubated on ice for 20 min and pelleted. The pellet was resuspended in 500 ul PBS and analyzed with flow cytometer (Becton Dickinson) at the HSCI-CRM Flow Cytometry Core Facility of MGH. The data were analyzed with FlowJo software.

### Tunicamycin stress test

EDEM3 KO and control HepG2 cells were treated with increasing amount of tunicamycin (Sigma-Aldrich) for 4 days (Graham et al., 2016). Cell viability was then analyzed with alamarBlue reagent (Fisher Scientific) according to the manufacturer’s protocol and absorbance was monitored at 570 nm.

### Plasma TG measurement and Western analysis of liver extracts

The mouse plasma TG was measured with triglyceride colorimetric assay kit (Cayman, 10010303) according to the manufacturer’s protocol. Liver lysates were prepared in 1XTBS containing 1% NP-40 and 0.1% doxycycline with the protease inhibitor cocktail (Roche). The liver lysate (∼50 μg total protein) was used for Western analysis with anti-EDEM3 and -LRP1 antibodies. All quantitation was normalized to the endogenous mouse GAPDH.

### Total RNA extraction and RNA-seq analysis

Total RNA from EDEM3 WT and KO HepG2 cells was extracted using RNeasy Mini Kit (Qiagen) according to the manufacturer’s procedures. RNA-seq analysis was performed at the Translational Genomics Core of Partners HealthCare.

### Total cell lipid extract preparation and metabolite mass spectrometry analysis

Preparation and metabolomics analysis of total cell lipid extracts from *EDEM3* KO and scramble control HepG2 cells were carried out as previously described (Mascanfroni et al., 2015). The blood lipid metabolite data of the individuals carrying SNP rs78444298 (P746S) were obtained from FHS 1000G GWAS cohorts as previously described.

### Statistic analysis

Statistic analyses for functional assays were performed using Prism 7 (GraphPad) with the student’s t-test or one way analysis of variance (ANOVA). Detailed group comparisons were described in individual figure legends. *p* values of ≤ 0.05 were considered to be statistically significant (*, *p*<0.05; **, *p* <0.01; ***, *p* <0.001).

## Acknowledgements

This work was supported by funding from National Heart, Lung and Blood Institute grant 5R33HL120781-05 to S.K. G.M.P. is supported by KO1HL125751, RO3HL141439, and RO1HL142711. We thank Partners Healthcare Personalized Medicine for the RNA-seq analysis, and Andrew Johnston for his contribution to the early stage of this project.

## Author contributions

Conceptulization, X.Y.X. and S.K.; Methodolody, X.Y.X., G.M.P., K.M., C.B.C., D.R. and S.K.; Formal analysis, X.Y.X., G.M.P., H.L., T.H.N., A.D., and K.B.; Investigation, X.Y.X., G.M.P., T.H.N., T.M., A.D., K.B.; Resources, Q.Y., R.S.V., and R.E.G.; Writing -Original draft, X.Y.X.; -editing, X.Y.X., G.M.P., T.H.N., T.M., A.D., H.L., R.S.V., and S.K.; -Supervision, X.Y.X. and S.K.; Funding acquisition, S.K.

## Supplemental figure legends

**sFig. 1. Deletion of *EDEM3* gene did not affect Dil-LDL uptake**. (**A**) Three individual EDEM3 scramble control (WT) and KO HepG2 (**A**) and Huh7 (**B**) clones were incubated with Dil-LDL. Tops, Representative images of the uptake assays from *EDEM3* KO and control cells. Bottoms, Quantification of the uptake assays from three clones of the control cells (>100 cells) and KO cells (>560 cells). Cellular Dil-VLDL particle areas were quantified using ImageJ. *p* values (t-test) were calculated by comparing the KO cells with the control cells. Bar, 25 mm.

**sFig. 2. Complementary analysis of *EDEM3* gene deletion in HepG2 cells with WT and P746S mutant EDEM3 on LRP1 expression. A.** The control and complementary HepG2 cells as in Fig. 7 were immunostained with antibodies against LRP1 and ApoB. The cells were analyzed under confocal microscope. **B**. Image quantification of LRP1 and ApoB signal was carried out using ImageJ from 100-200 cells per clone. *, p<0.05. *p* values (t-test) were calculated by comparing the KO+WT cells with the KO or KO+P746S cells. ANOVA *p* value<0.05 for LRP1 expression; *p* value<0.05 for ApoB signal.

**sFig. 3. Complementary analysis of *EDEM3* gene deletion in Huh7 cells with WT and P746S mutant EDEM3 on LRP1 expression.** Similar analysis as in sFig. 2 but for Huh7 cells (100-200 cells per clone). *p* values (t-test) were calculated by comparing the KO+WT cells with the KO or KO+P746S cells. *, p<0.05; **, p<0.01; ***, p<0.001. ANOVA *p* value<0.001 for LRP1 expression; *p* value<0.01 for ApoB signal.

**sFig. 4. Complementary analysis of *EDEM3* gene deletion in HepG2 cells with WT and P746S mutant EDEM3 on VLDL uptake. A.** The control and complementary HepG2 cells as in Fig. 7 were used for VLDL uptake assay as in Fig. 3. immunostained with antibodies against LRP1 and ApoB. The cells were analyzed under confocal microscope. **B**. Image quantification of LRP1 and ApoB signal was carried out using ImageJ from 100-269 cells per clone. *p* values (t-test) were calculated by comparing the KO+WT cells with the KO or KO+P746S cells. **, p<0.01. ANOVA *p* value<0.0001.

**sFig. 5. Complementary analysis of *EDEM3* gene deletion in Huh7 cells with WT and P746S mutant EDEM3 on VLDL uptake.** Similar analysis as in sFig. 4 but for Huh7 cells (100-220 cells for each clone). *p* values (t-test) were calculated by comparing the KO+WT cells with the KO or KO+P746S cells. *, p<0.05; **, p<0.01. ANOVA *p* value<0.001.

