## supplemental Table and figures for "EDEM3 modulates plasma triglyceride level through its regulation of LRP1 expression"

sTable 1: EDEM3 missense mutations associated with lipid phenotypes

| RSID | CHR:POS** | REF | ALT | Protein Change | Alternate Allele Frequency | Trait* | Sample Size | Beta*** | SE | p-value |
|---|---|---|---|---|---|---|---|---|---|---|
| rs141657255 | 1:184663288 | C | A | W903L | 0.03% | TG | 284295 | 0.0194 | 0.0803 | 0.8092 |
| | | | | | | LDL | 274616 | -0.0931 | 0.0809 | 0.2497 |
| | | | | | | HDL | 295092 | -0.0595 | 0.0806 | 0.4607 |
| rs9425343 | 184663537 | A | C | I820S | 39.15% | TG | 304422 | 0.0019 | 0.0027 | 0.4774 |
| | | | | | | LDL | 294565 | 0.0070 | 0.0027 | 0.0108 |
| | | | | | | HDL | 315135 | -0.0010 | 0.0027 | 0.7183 |
| **rs78444298** | **1:184672098** | **G** | **A** | **P746S** | **1.50%** | **TG** | **262076** | **-0.0502** | **0.0115** | **$1.2x10^{-05}$** |
| | | | | | | LDL | 253261 | 0.0005 | 0.0117 | 0.9645 |
| | | | | | | HDL | 272904 | 0.0045 | 0.0112 | 0.6838 |
| rs200489181 | 1:184681571 | T | C | N511S | 0.01% | TG | 283685 | 0.0451 | 0.1403 | 0.7478 |
| | | | | | | LDL | 273994 | -0.0428 | 0.1404 | 0.7603 |
| | | | | | | HDL | 294447 | 0.1643 | 0.1402 | 0.2413 |
| rs139183949 | 1:184690424 | G | A | T317I | 0.01% | TG | 284269 | 0.0697 | 0.1849 | 0.7062 |
| | | | | | | LDL | 274520 | -0.0814 | 0.1845 | 0.6592 |
| | | | | | | HDL | 294971 | -0.2787 | 0.1849 | 0.1317 |
| rs202039206 | 1:184695486 | C | T | R217Q | 0.71% | TG | 239249 | -0.0109 | 0.0177 | 0.5387 |
| | | | | | | LDL | 229888 | -0.0295 | 0.0182 | 0.1051 |
| | | | | | | HDL | 242531 | -0.0023 | 0.0177 | 0.8952 |
| rs201274616 | 1:184703729 | T | C | R132G | 0.02% | TG | 274028 | -0.0297 | 0.1040 | 0.7749 |
| | | | | | | LDL | 264439 | -0.0767 | 0.1063 | 0.4705 |
| | | | | | | HDL | 284805 | 0.0101 | 0.1051 | 0.9236 |

*TG was log transformed before analysis
**Positions are on hg19
***Beta is in standard deviation units

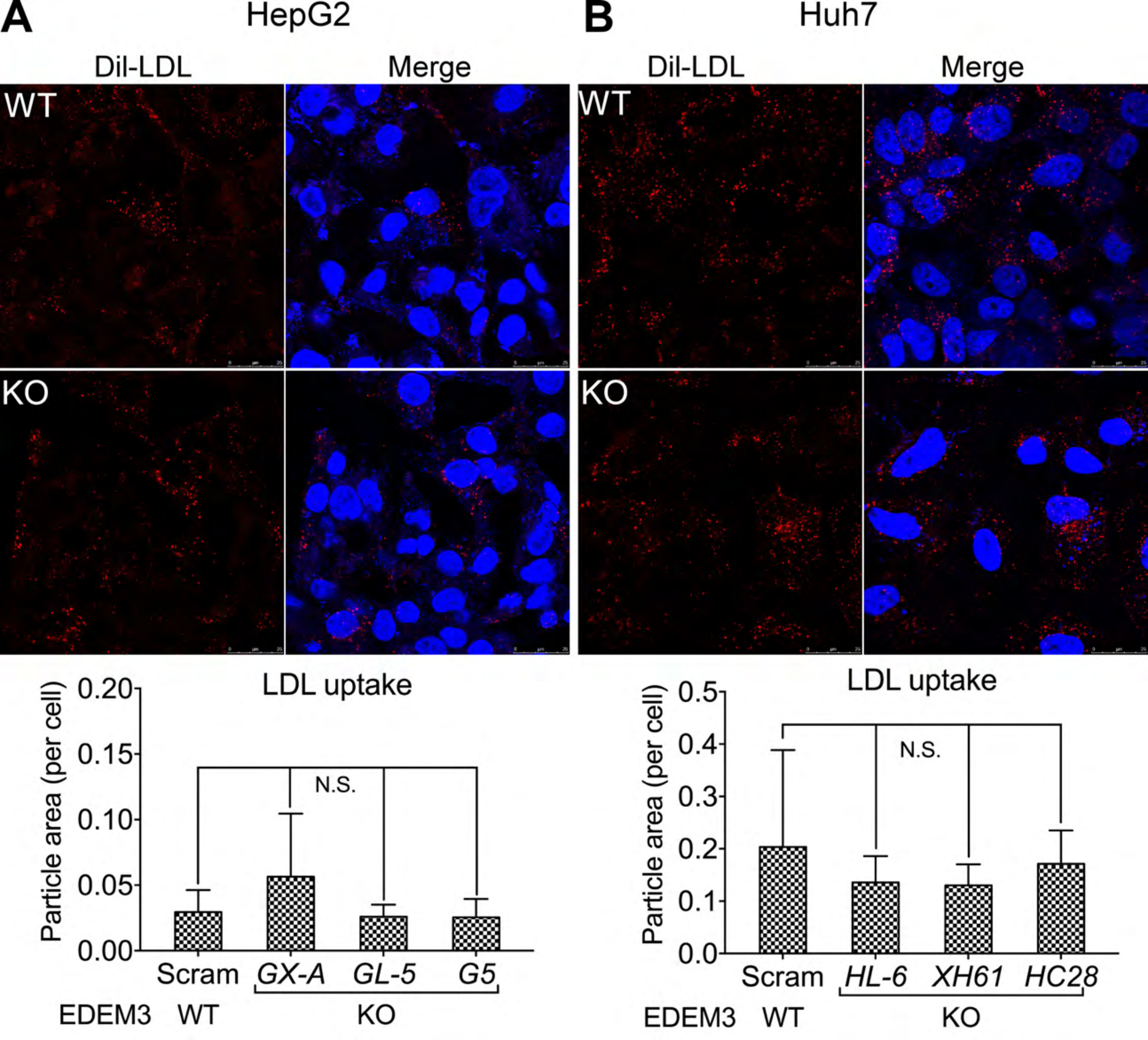

A
HepG2
Dil-LDL
Merge
WT
KO
LDL uptake
Particle area (per cell)
0.20
0.15
0.10
0.05
0.00
N.S.
Scram
GX-A
GL-5
G5
EDEM3
WT
KO
B
Huh7
Dil-LDL
Merge
WT
KO
LDL uptake
Particle area (per cell)
0.5
0.4
0.3
0.2
0.1
0.0
N.S.
Scram
HL-6
XH61
HC28
EDEM3
WT
KO

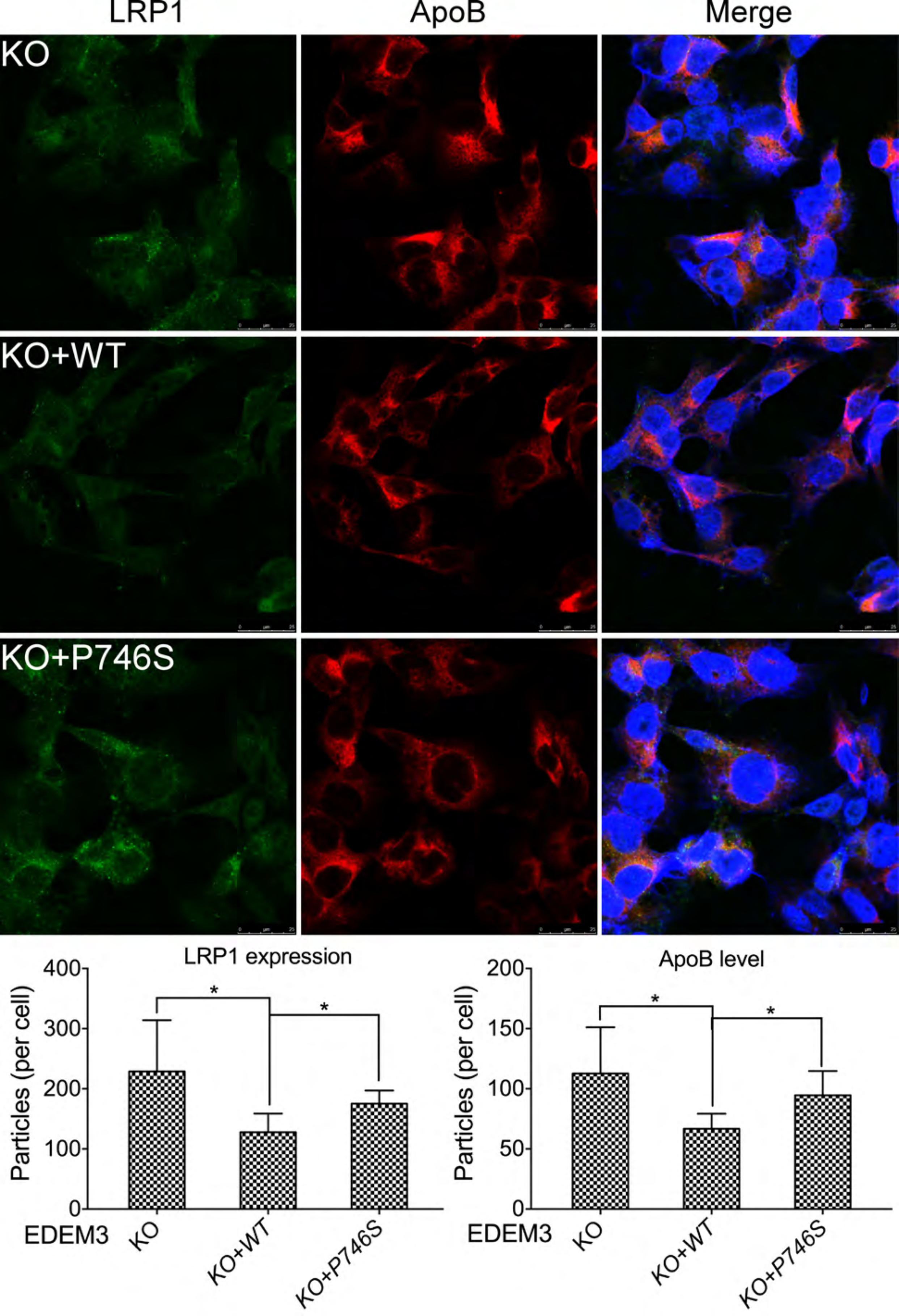

LRP1
ApoB
Merge
KO
KO+WT
KO+P746S
LRP1 expression
Particles (per cell)
400
300
200
100
0
*
*
EDEM3
KO
KO+WT
KO+P746S
ApoB level
Particles (per cell)
200
150
100
50
0
*
*
EDEM3
KO
KO+WT
KO+P746S

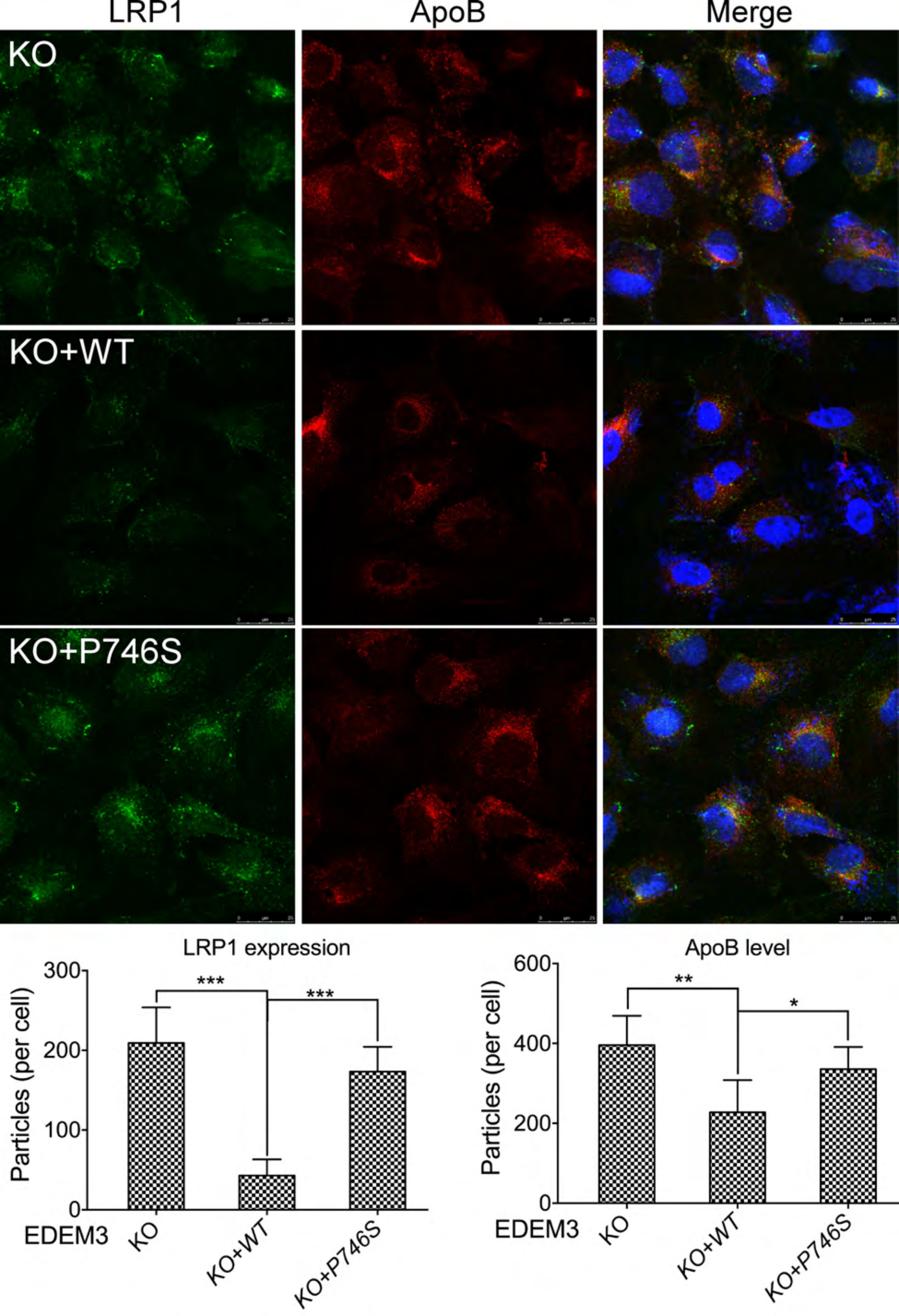

LRP1
ApoB
Merge
KO
KO+WT
KO+P746S
LRP1 expression
Particles (per cell)
300
200
100
0
***
***
EDEM3
KO
KO+WT
KO+P746S
ApoB level
Particles (per cell)
600
400
200
0
**
*
EDEM3
KO
KO+WT
KO+P746S

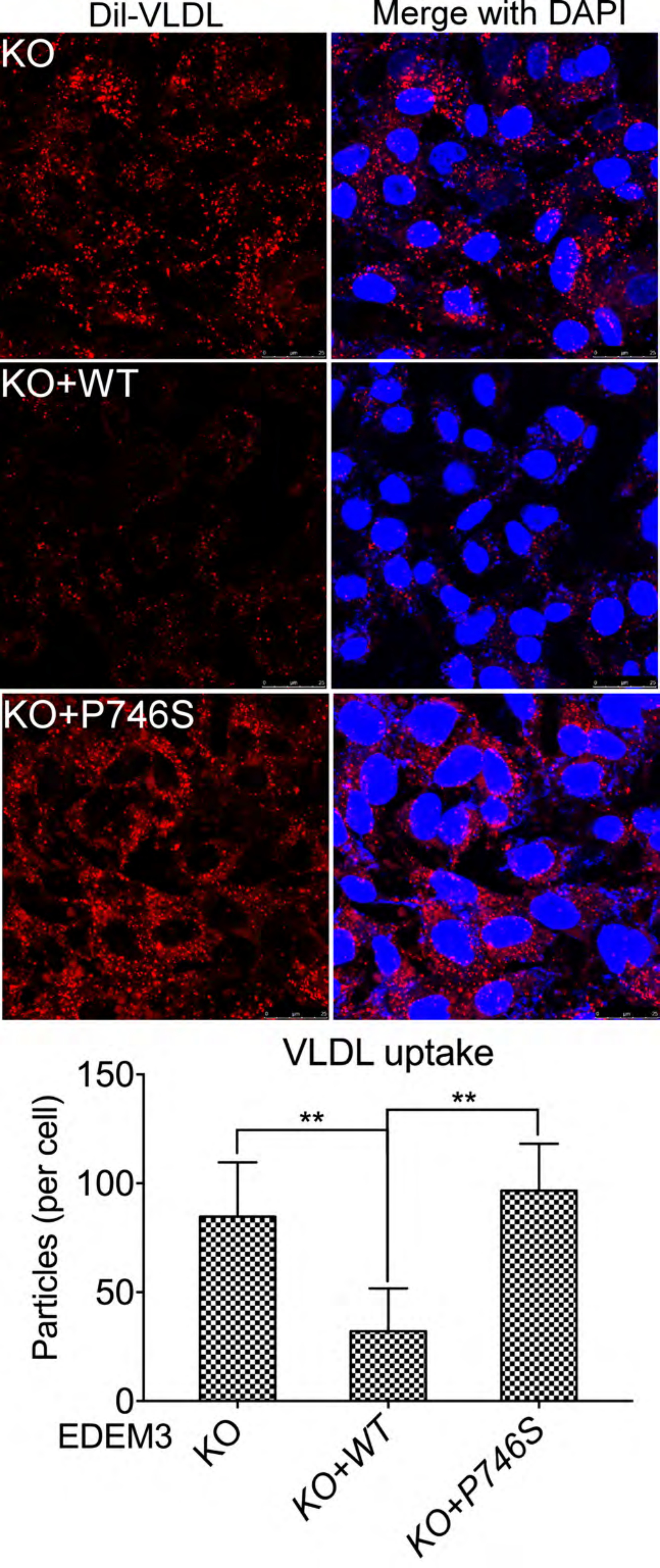
DiI-VLDL
Merge with DAPI
KO
KO+WT
KO+P746S
VLDL uptake
Particles (per cell)
150
100
50
0
**
**
EDEM3
KO
KO+WT
KO+P746S

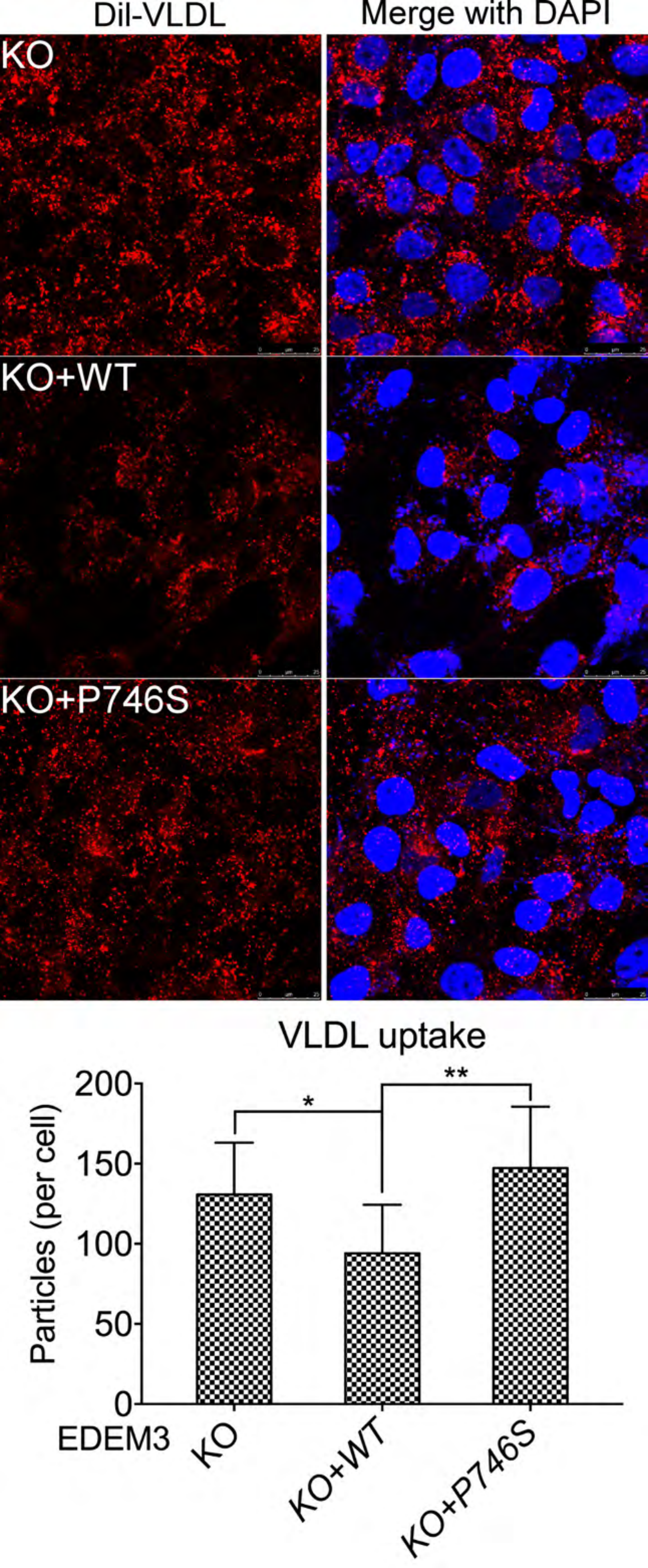

Dil-VLDL
Merge with DAPI
KO
KO+WT
KO+P746S
VLDL uptake
Particles (per cell)
200
150
100
50
0
*
**
EDEM3
KO
KO+WT
KO+P746S
